# Sestrins regulate age-induced deterioration of muscle stem cell homeostasis

**DOI:** 10.1101/2020.12.15.422905

**Authors:** Benjamin A. Yang, Jesus Castor-Macias, Paula Fraczek, Lemuel A. Brown, Myungjin Kim, Susan V. Brooks, Jun Hee Lee, Carlos A. Aguilar

## Abstract

The health and homeostasis of skeletal muscle is preserved by a population of tissue resident stem cells called satellite cells. Young healthy satellite cells maintain a state of quiescence, but aging or metabolic insults results in reduced capacity to prevent premature activation and stem cell exhaustion. As such, understanding genes and pathways that protect satellite cell maintenance of quiescence are needed. Sestrins are a class of stress-inducible proteins that act as antioxidants and inhibit the activation of the mammalian target of rapamycin complex 1 (mTORC1) signaling complex. Despite these pivotal roles, the role of Sestrins has not been explored in adult stem cells. Herein, we show that Sestrin1,2 loss results in hyperactivation of the mTORC1 complex, increased propensity to enter the cell cycle and shifts in metabolic flux. Aging of Sestrin1,2 knockout mice demonstrated a loss of MuSCs and reduced ability to regenerate. These findings demonstrate Sestrins function to help maintain MuSC metabolism that supports quiescence and against aging.

**Highlights:**

- Sestrin deficiency alters mTORC1 signaling in muscle stem cells (MuSCs).
- In young mice, Sestrins are dispensable for regenerative responses of MuSCs.
- Sestrin deficiency accelerates age-dependent loss and dysfunction of MuSCs.

## Introduction

Tissue resident stem cells are crucial regulators of homeostasis that resist perpetual activation through mitotic quiescence, a reversible restraint of entry into the G0 phase of the cell cycle. After injury, tissue-specific stem cells leave quiescence, activate, and proliferate, generating either committed progenitors that differentiate to repair tissue or return towards quiescence to replenish the stem cell pool^1^. An excellent example of this process is in skeletal muscle, a postmitotic tissue supported by Pax7^+^ muscle stem cells (MuSCs)^2^, also known as satellite cells, that reside in a quiescent niche between the basal lamina and sarcolemma. Upon activation, MuSCs migrate along myofibers and asymmetrically produce myogenic progenitors (myoblasts) that fuse with themselves and existing myofibers to regenerate and repair damaged tissue^3^. MuSC quiescence is critically determined both by internal mechanisms such as chromatin state^4,5^ and external communication with the extracellular matrix (ECM)^6^ and other cell types^7^. Defects in the balance between quiescence and activation have been observed in cachexia^8^, sarcopenia^9^, and in diseases such as Duchenne muscular dystrophy (DMD)^7,10^, highlighting a clinical need to understand the molecular mechanisms by which healthy MuSCs balance quiescence and proliferative cues.

A critical yet underexplored element of quiescence regulation is the maintenance of metabolic homeostasis. Quiescent MuSCs utilize fatty acid oxidation (FAO) in quiescence but stressors such as reactive oxygen species (ROS), DNA damage^1,11^, and inflammation, shift metabolism away from FAO towards glycolysis, which in turn prompts MuSCs to activate and leave the quiescent state^12^. Several genes influence quiescence and FAO, including the forkhead box O (FOXO) family of transcription factors, which maintains bioenergetic demands^13^ and nutrient consumption in MuSCs by inhibiting oxidative stress and activation of the mammalian target of rapamycin complex (mTORC) signaling through the Phosphatidylinositide 3 kinase (PI3K)/AKT pathway ^14–17^. Age-induced deterioration of quiescence maintenance and hyperactivation of both the PI3k/AKT and mTORC pathways leads to increased stem cell turnover and impaired self-renewal^9,18,19^. Inhibitors of mTORC activity^20^ such as the tuberous sclerosis proteins 1 and 2 (TSC1/2) complex^21^, and the AMP activated protein kinase (AMPK) complex^22^, have both been shown to regulate MuSC metabolism and longevity. Another mTORC inhibitor called Sestrins^23^, a set of conserved proteins encoded by Sesn1, Sesn2, and Sesn3 in mammals, mediate metabolic responses to oxidative stress by physically activating AMPK signaling and indirectly inhibiting mTORC 1 pathways^24–26^. In skeletal muscle, Sestrin-mediated inhibition of mTORC 1 and AKT pathways upregulates autophagy and downregulates anabolic pathways to maintain proteostasis and organelle quality, ultimately preserving muscle mass and force^27^. While Sestrins are implicated in muscle pathophysiology and regulate mTORC1 and aging, their role in MuSCs and stem cells in general remains unexplored.

Herein, we utilized Sestrin1,2 knockout (SKO) murine models to evaluate the influence of Sestrins on MuSC functions. Muscles of SKO mice resulted in hyperactivation of the mTORC1 complex in MuSCs, even in the absence of regenerative stimuli. This led to an increased propensity to enter the cell cycle and accumulation of ROS. Despite the alterations in basal stem cell status, application of a muscle injury to SKO mice revealed highly similar regenerative trajectories as wild-type matched controls. We used RNA-sequencing and metabolic flux modeling to show that SKO MuSCs displayed modifications to FAO and other amino acid metabolic fluxes, which impinge on mTORC1 and reduce ability to maintain quiescence. To probe deeper into this result, we aged SKO mice and observed an accelerated age-dependent loss of MuSCs. Repeated muscle injuries to aged SKO mice showed increases in fibrosis and smaller cross-sectional areas of myofibers indicating reductions in regenerative potential. These studies demonstrate a unique role for Sestrins as arbiters of MuSC metabolism and protection against aging.

## Results

### Sestrin Loss Upregulates mTORC1 in Unstimulated Muscle Stem Cells

To examine the role that Sestrins play in regulating muscle stem cells (MuSC), we isolated hindlimb muscles from 2-month old uninjured C57BL6J (wild-type or WT) and SKO mice. Comparisons of myofiber size and fiber typing revealed no significant differences for SKO muscles (Supp. Fig. 1). No statistically significant change in the total number of Pax7^+^ MuSCs was observed between WT and SKO quadriceps samples (Fig. 1a-b). However, in situ staining of Pax7 and the phosphorylated form of ribosomal protein S6 (pS6), an mTORC 1 target, in quadriceps muscle revealed a substantial increase in the fraction of pS6^+^ / Pax7^+^ MuSCs for SKO muscles compared to WT controls (Fig. 1a-b). The increase in pS6 fraction and concomitant hyperactivation of mTORC 1 in SKO MuSCs are in line with previous studies of TSC1 knockout^21^, showing that Sestrins are negative regulators of mTORC 1 and suggest reduced quiescence in SKO MuSCs.

**Figure 1.**
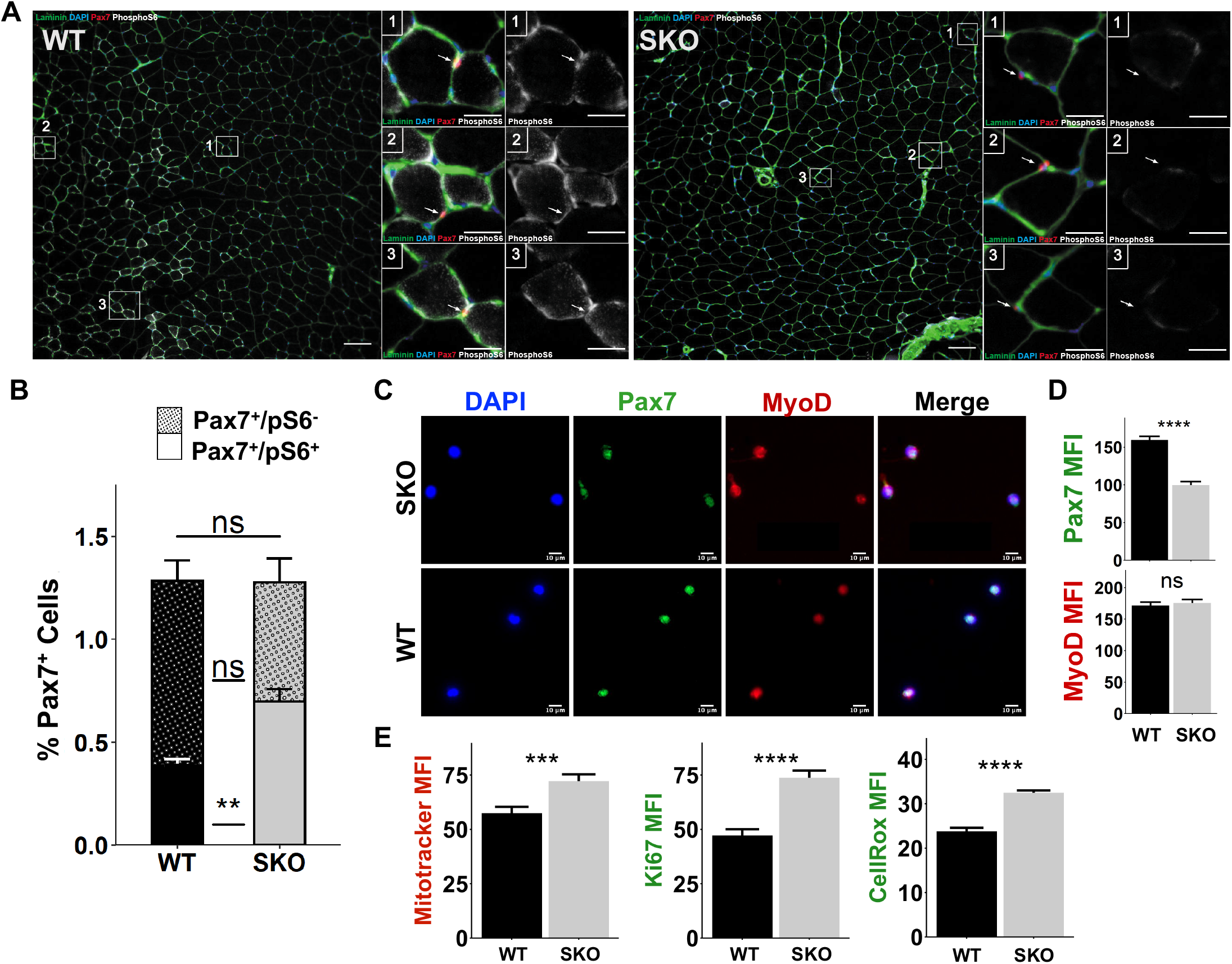
Loss of Sestrin1,2 induces hyper-activation of mTORC1 signaling in muscle stem cells. **A)** Representative in situ immunohistochemical images of whole quadriceps muscle sections from wild type (WT, left) and Sestrin1,2^-/-^ (SKO, right) mice stained for DAPI (blue), Pax7 (red), Laminin (green), and PhosphoS6 (pS6, white), a marker of mTORC1 activation. Scale bars are 100 μm. Magnified images show Pax7^+^ cells for each condition. Scale bars are 25 μm. **B)** Quantification of pS6^+^ (stippled) and pS6^-^ (plain) Pax7^+^ cells as a percentage of all nuclei in whole WT and SKO quadriceps sections. Statistical comparisons are two-sided Mann-Whitney *U*-tests. (WT: n=3, SKO: n=3). **C)** Immunofluorescence staining for Pax7 (green), MyoD (red) and DAPI (blue) in SKO and WT MuSCs fixed immediately after isolation. Scale bars are 10 μm. **D)** Mean fluorescence intensity (MFI) of Pax7 and MyoD in SKO and WT MuSCs fixed immediately after isolation. Statistical comparisons are two-sided unpaired Student’s *t*-tests. **E)** Quantification of MFI from Mitotracker, Ki67, and CellRox in WT and SKO MuSCs fixed immediately after isolation. Statistical comparisons are two-sided unpaired Student’s *t*-tests. All data are shown as mean ±SEM. **p<0.01, ***p<0.001, ****p<0.0001.

### Sestrin Loss Reduces Ability to Maintain MuSC Quiescence

To probe further if Sestrin loss and associated increases in mTORC 1 signaling influenced MuSCs, WT and SKO hind limb muscles (tibialis anterior, extensor digitorum longus, gastrocnemius, and quadriceps) were extracted and MuSCs isolated using fluorescent activated cell sorting^28^ (FACS) with both negative (Sca-1^-^, CD45^-^, Mac-1^-^, Ter-119^-^) and positive surface markers (CXCR4^+^, β1-integrin^+^) (Supp. Fig. 2a). Freshly isolated SKO MuSCs displayed decreased Pax7 expression with higher levels of ROS (CellROX) and mitochondria (Mitotracker) compared to WT MuSCs (Figs. 1c-e, Supp. Fig. 2f-g). SKO MuSCs also displayed increases in proliferation (as measured by Ki67) but similar levels of MyoD, suggesting stronger resistance to activation in WT than SKO MuSCs (Figs. 1d-e, Supp. Fig. 2f). Culture of both types of MuSCs in activating conditions for 72 hours showed similar levels of Pax7, indicating comparable levels of self-renewal capacity, but increased MyoD expression in SKO MuSCs (Supp. Fig. 3a-b). Given that autophagy is critical for MuSC activation^18^ and that Sestrins suppress autophagic degradation^24^, we performed immunofluorescence (IF) staining of p62/sequestosome (SQSTM) at the same time points. We observed no differences between WT and SKO MuSCs (Supp. Fig. 2b,d, Supp. Fig. 3c-d), and no differences in total AMPKα1 levels between WT and SKO MuSCs were observed at either timepoint (Supp. Fig. 2c,e, Supp. Fig. 4e-f). Integrating these results suggests that Sestrins influence MuSCs in homeostasis but do not significantly affect catabolic activity or activation dynamics.

### Sestrin Deficiency Does Not Alter Injury Responses of Muscle Stem Cells in Young Mice

To glean deeper insights into the consequences of alterations in homeostasis from Sestrin1,2 knockout, WT and SKO hind limb muscles (tibialis anterior, gastrocnemius, and quadriceps) were injured through BaCl_2_ injections. MuSCs were isolated with FACS before and after injury (0, 7, and 21 days post injury – dpi), and submitted for RNA-Sequencing^29^ (RNA-Seq) (Fig. 2a). Gene expression profiles from WT and SKO samples demonstrated strong agreement between biological replicates at each time point (Spearman > 0.97) (Supp. Fig. 4a) and principal component analysis (PCA) revealed that while WT and SKO MuSCs were transcriptionally dissimilar before injury, they migrated along similar regeneration trajectories (Fig. 2b). A total of 982 differentially expressed genes were identified across all time points, the majority of which (~90%) were uniquely expressed at 0 dpi (Fig. 2c, Supp. Fig. 4b). Dirichlet Process Gaussian Process (DPGP) clustering revealed time-based differences in enriched GO terms and KEGG pathways (Fig. 2d, Supp. Fig. 4c-e). Visualizing the DPGP clustered genes in PCA space showed that the temporal dynamics of clusters 1, 2, 4, and 7 were predominantly driven by genes that were differentially expressed as a result of injury (Supp. Fig. 4c). Consistent with WT MuSCs displaying variations in metabolism that contribute to enhanced quiescence, genes in cluster 1 that were upregulated in WT before injury were enriched for terms related to fatty acid derivative metabolism (e.g. Cyp2s1, Fabp5) and restraint of activation (e.g. Mstn, Snai2). Cluster 7 was enriched for genes comprising cell-cycle regulation terms (e.g. Cdk1, Ccna2, Ccnb1, Cdc25c) and summing these results further suggest that SKO MuSCs display alterations in metabolism and reductions in the ability to retain quiescence. To probe further into changes in homeostasis, we evaluated differentially expressed genes at 0 dpi. This analysis revealed that SKO MuSCs contained upregulated genes for cell cycle checkpoints and oxidative stress metabolism while WT MuSCs upregulated genes related to ribosomal activity, and inhibition of muscle tissue development (Fig. 2e). Given the variation in metabolic genes between SKO and WT MuSCs, we used genome-scale metabolic modeling to assess the relationship between 3,744 metabolic reactions, 2,766 metabolites, 1,496 metabolic genes and 2,004 metabolic enzymes^5,30^. The model predicted that uninjured SKO MuSCs upregulated metabolic flux through acetyl CoA synthetase, and glycine, serine, and threonine metabolism (Supp. Fig. 5). Given that mTORC1 is regulated by acetyl CoA^31^, these results are consistent with over-activated mTORC1 in SKO MuSCs and increases in expression of acetyl-transferases and deacetylases (Supp. Fig. 4f). Collectively, the observed changes in expression show that knocking out Sesn1,2 does not significantly alter muscle regeneration but rather drives alterations in metabolism that contribute to reduced quiescence in MuSCs.

**Figure 2.**
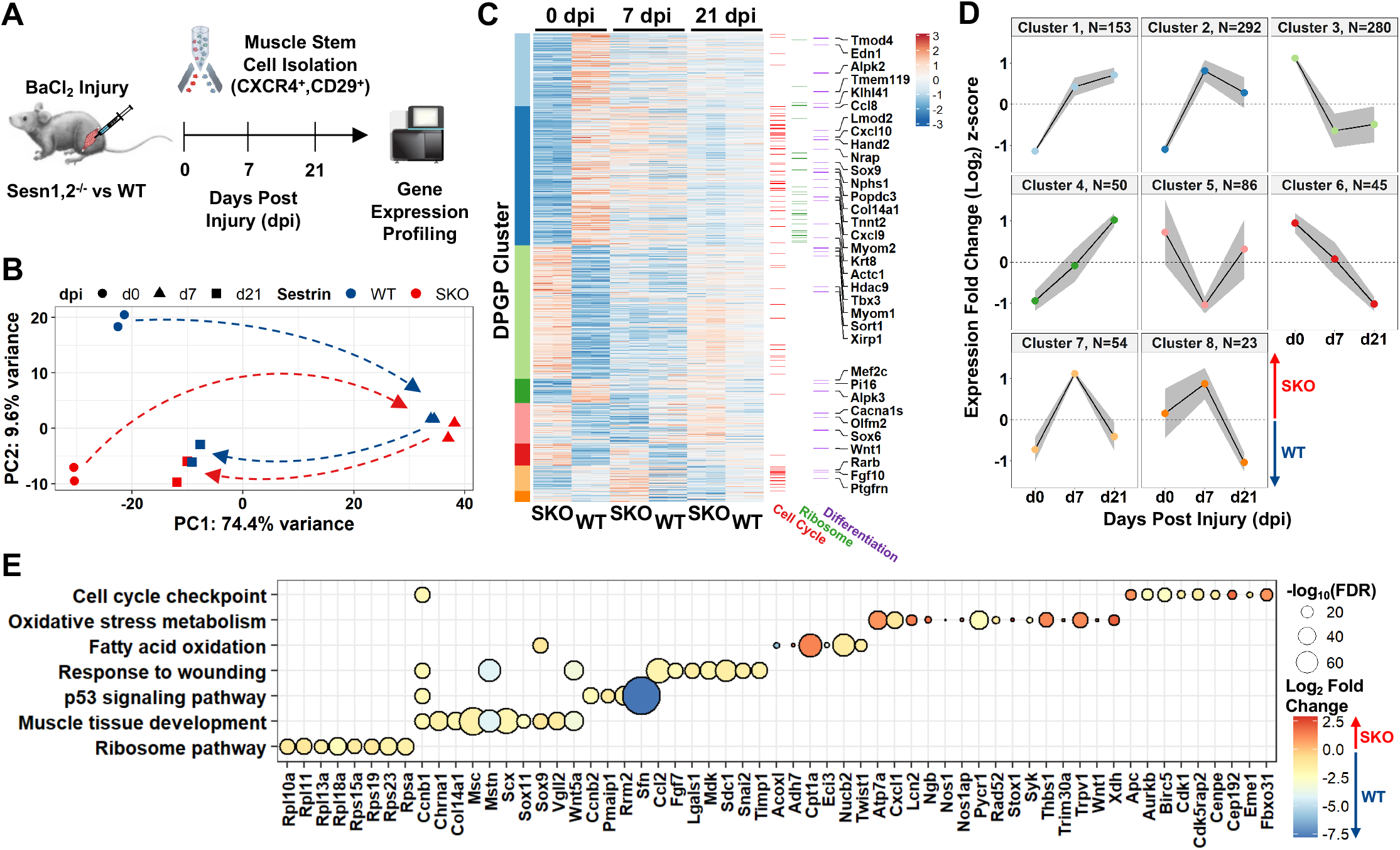
Sesn1/2 Knockout muscle stem cells show robust regenerative responses despite having strongly altered basal-level transcriptomes. **A)** Schematic diagram of BaCl_2_ injury model in the hindlimbs of WT and SKO mice followed by muscle stem cell (MuSC) isolation via FACS and RNA-Seq profiling before and during regeneration. **B)** Principal component analysis (PCA) of MuSC replicates from WT and SKO mice. **C)** Standardized heatmap (z-scores) of 982 differentially expressed genes (SKO over WT) from WT and SKO MuSCs grouped by Dirichlet Process Gaussian Process (DPGP) clusters. Genes that participate in the cell-cycle (Buettner et al., 2015), the Ribosome KEGG pathway (mmu03010), and the muscle cell differentiation GO term (GO:0042692) are marked on the right. **D)** DPGP mixture model-based clustering of mean gene expression time series z-scores (gray bars are 2x standard deviation and black line is cluster mean). **E)** Gene set terms and KEGG pathways at 0 days post injury (dpi) plotted against constituent genes.

### Aging Coupled with Sestrin Loss Decreases Muscle Stem Cell Number and Function

One way to evaluate changes in quiescence and long-term homeostasis of MuSCs is through natural aging, whereby increases in oxidative damage and negative changes in proteostasis promote constitutive MuSC activation and exhaustion^9,18^. We therefore examined gene expression datasets obtained from young and aged MuSCs before and after injury^5^ and observed aged MuSCs exhibit reductions in Sestrin1 expression during regeneration, which correlated with previous observations of reduced inability to maintain quiescence (Supp. Fig. 6a). To probe deeper into these observations, we aged SKO and WT mice to middle age (14 months), when TSC1,2 knockout demonstrate accelerated aging phenotypes^32^. We extracted hind limb muscles from middle-aged SKO and WT mice and observed slightly smaller myofibers and comparable numbers of centrally nucleated fibers, fiber types and p62 levels for SKO muscles compared to WT (Fig. 3a-c, Supp. Fig. 6b-d,e,g). Enumeration of Pax7^+^ MuSCs revealed a significant decrease in the overall number and an increase in the pS6 fraction of MuSCs for SKO compared to WT (Fig. 3d-e). These results are consistent with previous reports of MuSC loss in TSC1,2 knockout mice, and their inability to maintain quiescence^21^.

**Figure 3.**
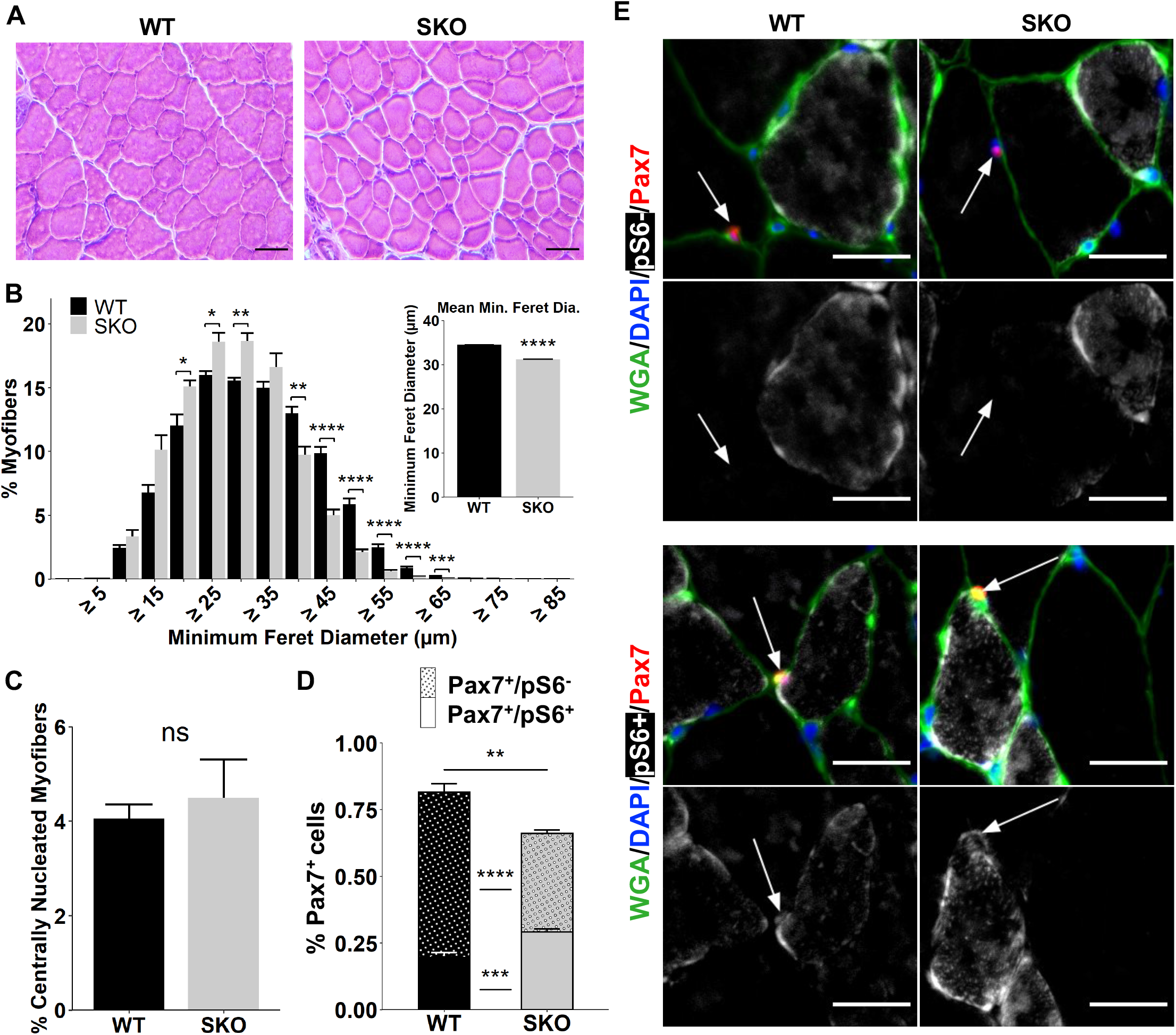
Sustained mTORC1 activation from loss of Sestrin1,2 in aging induces loss of muscle stem cells. **A)** Representative images of Hematoxylin & Eosin (H&E) staining. Scale bars are 100 μm. **B)** Size distributions of WT and SKO myofibers. Inset shows mean size per condition. Statistical comparisons are two-sided unpaired Student’s *t*-tests with Holm multiple testing correction for the histogram. (WT: n=5, SKO: n=3). **C)** Percentage of centrally nucleated myofibers in whole uninjured TA sections from WT and SKO mice. Statistical comparison is Mann-Whitney *U*-test. (WT: n=5, SKO: n=3). **D)** Quantification of pS6^+^ (stippled) and pS6^-^ (plain) Pax7^+^ cells as a percentage of all nuclei in whole uninjured TA muscle sections. Statistical comparisons are twosided Mann-Whitney *U*-tests. (WT: n=3, SKO: n=3). **E)** Representative immunohistochemical images showing pS6^-^ (top) and pS6^+^ (bottom) MuSCs in whole uninjured TA muscle sections from WT and SKO mice. Sections were stained with antibodies against Pax7 (red) and pS6 (white), and counterstained with DAPI (blue). Connective tissue was stained with wheat germ agglutinin (WGA) (green). Arrows mark Pax7^+^ cells. Scale bars are 25 μm. All data are shown as mean ± SEM. *p<0.05, **p<0.01, ***p<0.001, and ****p<0.0001. Unmarked comparisons lack statistical significance.

To further examine if changes in regenerative capacity of MuSCs are modified with loss of Sestrins during aging, we performed repetitive injuries by injecting BaCl_2_ into the TA muscle (injuries spaced 21 days apart between injections) (Fig. 4a). SKO muscles displayed reductions in fiber size after repeated injuries (Fig. 4b-c), increases in the number of centrally located nuclei (Fig. 4d), enhanced collagen deposition (Fig. 4e-f), and loss of type IIx and gain of type IIb fiber types (Supp. Fig. 6f,h). The total number of MuSCs was also slightly reduced in SKO relative to WT (Fig. 4g-h), with no change in p62 levels (Supp. Fig. 6b-c). Summing these results demonstrates that Sestrin deficiency produces an attenuated regenerative response owing to a reduced MuSC population akin to aging.

**Figure 4.**
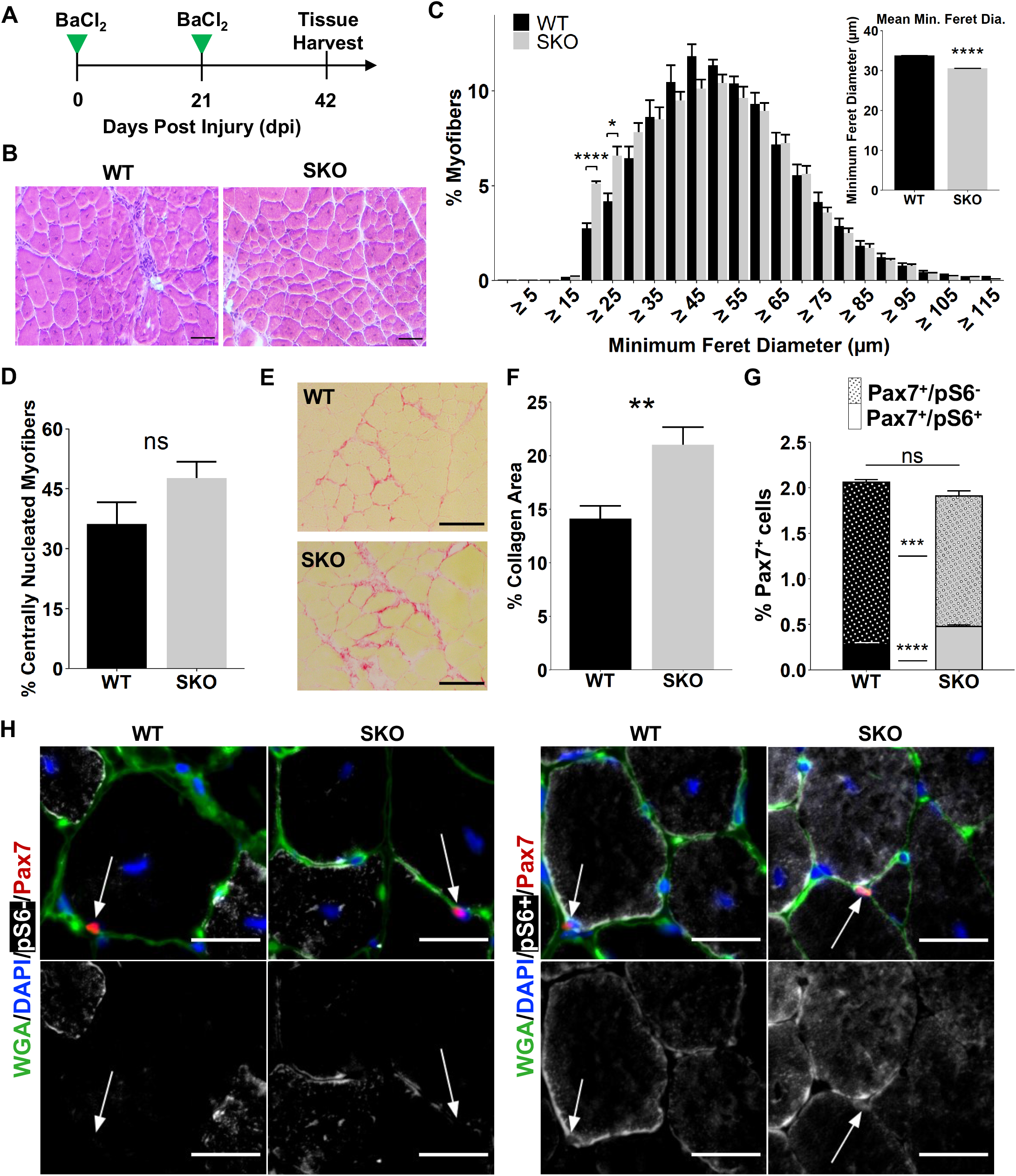
Loss of Sestrin1,2 in aging muscle impairs regeneration and promotes muscle stem cell loss following injury. **A)** Schematic of BaCl_2_ double injury model. **B)** Representative images of Hematoxylin & Eosin (H&E) staining. Scale bars are 100 μm **C)** Size distribution of WT and SKO myofibers. Statistical comparisons are two-sided unpaired Student’s *t*-tests with Holm multiple testing correction for the histogram. Inset is mean myofiber size per condition. (WT: n=4, SKO: n=3). **D)** Percentage of centrally nucleated myofibers in whole injured WT and SKO TA muscle sections. (WT: n=4, SKO: n=3). Statistical comparison is two-sided Mann-Whitney *U*-test. **E)** Representative images of Picrosirius red-stained collagen in whole injured TA muscle sections. Muscle fibers are shown in yellow. Scale bars are 100 μm. **F)** Collagen area fraction in whole TA muscle sections as determined by Picrosirius red staining. Statistical comparison is two-sided unpaired Student’s *t*-test. (WT: n=3, SKO: n=4). **G)** Quantification of pS6^+^ (stippled) and pS6^-^ (plain) Pax7^+^ cells as a percentage of all nuclei in whole injured TA muscle sections.

## Discussion

Tissue homeostasis is critically dependent on the maintenance and activity of stem cells, which in turn are regulated by metabolic status. A focal point of metabolic signaling is the nutrient-sensitive protein complex mTORC1, which balances protein biosynthesis and autophagy to meet anabolic demands^33^. Regulation of mTORC1^34^ is essential to prevent stem cell over-activation^35^ and maintain quiescence^36^ and loss of mTORC1 inhibition results in depletion of hematopoietic stem cells^37^. Sestrins are a unique class of metabolic regulators that titrate mTORC1 signaling and act as oxidative stress sensors, but their roles in adult stem cells has yet to be evaluated. Our results show that Sestrins contribute to MuSC homeostasis by inhibiting mTORC1 and maintaining metabolism supportive of quiescence. The loss of Sestrins resulted in upregulation of genes related to oxidative stress metabolism and a reduced capacity to resist activation. Persistent increases in oxidative stress from hyperactivation of mTORC1 has been shown to promote an aged phenotype in muscle^38^ and prime MuSCs for activation through accelerated cell cycle entry and increased mitochondrial activity^21^. Our studies are consistent with these observations and demonstrate that Sestrins fine tune basal metabolism of MuSCs via stress-inducible genes with antioxidant functions such as nuclear factor (erythroid-derived 2)-like 2 (Nrf2)^39^ and other ROS-sensitive factors such as p53 and the FOXO family^24^. Given hyperactivated mTORC1 promotes mitochondrial biogenesis and oxidative phosphorylation, a source of ROS generation and electron leakage^40^, our results suggest Sestrins balance mTORC1 and the cellular redox state in MuSCs.

The regeneration of muscle is mediated through MuSC actions but in aging, the ability of MuSCs to repair tissue and self-renew declines^41^. This is in part owing to accumulation of DNA damage^42^, metabolic reprogramming in response to increases in oxidative stress^43^, and altered chromatin packaging^5^. Our results demonstrate that Sestrin-deficient muscle displayed comparable regenerative trajectory in young muscle (2-3 mos), but exhibit pathological aging in middle age (14 mos) through MuSC loss compounded with impaired regenerative capacity, independent of changes in autophagic flux. These results may be driven by alterations in mitochondria and production of energy. Previously, we observed that Sestrin loss resulted in attenuation of mitochondrial biogenesis and maximal respiratory capacity. Both are pivotal for proper muscle repair via MuSCs and interruptions to mitochondrial generation and bioenergetics can be detrimental for MuSC differentiation. We posit that Sestrins provide a control layer for MuSC mitochondria to protect against aging insults, but this control layer is diminished in aging and results in imbalances of ROS levels^44^. In support of this, both Sestrin1 and 2 exhibited decreases in expression in aged MuSCs compared to young^5^.

Integrating our data suggest Sestrins are important regulators of MuSC metabolism and the quiescent state. Several studies have shown that hyperactivation of mTORC1 results in stem cell loss^34,36^ and interruptions to the maintenance of quiescence but none have yet demonstrated this behavior through Sestrins. Accordingly, Sestrin-dependent changes may be strong targets for restoring MuSC metabolism in aging and expand our understanding of metabolic regulation of stem cells across lifespan.

## Acknowledgments

The authors thank Kanishka de Silva, and the University of Michigan DNA Sequencing Core for assistance with sequencing. The authors also thank Anna Shcherbina for insights into bioinformatics analysis, and members of the Aguilar, Lee and Brooks laboratories. Research reported in this publication was partially supported by the National Institute of Arthritis and Musculoskeletal and Skin Diseases of the National Institutes of Health under Award Number P30 AR069620 (CAA, SVB), the 3M Foundation (CAA), American Federation for Aging Research Grant for Junior Faculty (CAA), the Department of Defense and Congressionally Directed Medical Research Program W81XWH2010336 (CAA), the University of Michigan Geriatrics Center and National Institute of Aging under award number P30 AG024824 (CAA, SVB), the University of Michigan Biomedical Engineering Department (CAA), National Institute on Aging P01 AG051442 (SVB), and the National Institute for Biomedical Imaging and Bioengineering Training Award T32 EB005582 (BAY). The content is solely the responsibility of the authors and does not necessarily represent the official views of the National Institutes of Health.

## Accession Code

### Author contributions

B.A.Y., P.F., J.C.M., L.A.B., and M.K. performed experiments. B.A.Y., and C.A.A. analyzed data. S.V.B., J.H.L., and C.A.A. designed the experiments. B.A.Y., and C.A.A. wrote the manuscript with additions from other authors.

### Materials & Methods

| REAGENT or RESOURCE | SOURCE | IDENTIFIER |
| --- | --- | --- |
| **Antibodies** |  |  |
| Mouse anti-human myosin (embryonic), Isotype: Mouse IgG1, supernatant | Developmental Studies Hybridoma Bank | F1.652s |
| Goat anti-rabbit H+L, Alexa Fluor 555 conjugate | Invitrogen | A21428 |
| APC anti-mouse Ly-6A/E (Sca-1), clone: D7, isotype: Rat IgG2a, $\kappa$ | BioLegend | 108112 |
| APC anti-mouse CD45, clone: 30-F11, isotype: Rat IgG2b, $\kappa$ | BioLegend | 103112 |
| APC anti-mouse/human CD11b, clone: M1/70. Isotype: Rat IgG2b, $\kappa$ | BioLegend | 101212 |
| APC anti-mouse TER-119, clone: TER-119, isotype: Rat IgG2b, $\kappa$ | BioLegend | 116212 |
| PE anti-mouse/rat CD29, clone: HMβ1-1, isotype: Armenian Hamster IgG | BioLegend | 102208 |
| Biotin Rat Anti-Mouse CD184, clone: 2B11/CXCR4 (RUO), isotype: Rat IgG2b, $\kappa$, lot # 6336587 | BD Bioscience | 551968 |
| Streptavidin PE-Cyanine7, lot # 4290713 | eBioscience | 25-4317-82 |
| 4',6-Diamidino-2-Phenylindole, Dihydrochloride (DAPI), FluoroPure™ grade | Invitrogen | D21490 |
| M.O.M. mouse IgG blocking reagent | Vector Laboratories | BMK-2202 |
| BA-D5 IgG2b | DSHB |  |
| SC-71 | DSHB |  |
| BF-F3 | DSHB |  |
| Alexa Fluor 647 Goat anti Mouse IgG2b | Invitrogen | A-21242 |
| Alexa Fluor 488 Goat anti Mouse IgG1 | Invitrogen | A-21121 |
| Alexa Fluor 555 Goat anti Mouse IgM | Invitrogen | A-21426 |
| WGA 405 | Biotium | 29027 |
| WGA 488 | Biotium | 29022 |
| Tyramide SuperBoost Goat anti Mouse AF555 kit | Invitrogen | B40913 |
| Tyramide SuperBoost Goat anti Mouse AF488 | Invitrogen | B40912 |
| Pax7 from supernatant | DSHB University of Iowa |  |
| anti-rabbit Phospho-S6 | Cell Signaling | 4858 |
| Alexa Fluor 647 Goat anti rabbit | Invitrogen | A-21244 |
| Alexa fluor 555 Goat anti rabbit | Invitrogen | A-21428 |
| SQSTM1/p62 Rabbit | Cell signal | 23214 |
| **Chemicals, Peptides, and Recombinant Proteins** |  |  |
| DMEM/F-12, HEPES | Gibco | 11330032 |
| DMEM, high glucose, pyruvate | Life Technologies | 11995065 |
| Ham's F-10 Nutrient Mix | Life Technologies | 11550043 |
| Gentamycin/Amphotericin-B (GA-1000) | Lonza | CC-4083 |
| Dexamethasone | G-Biosciences | API-04 |
| Human Epidermal Growth Factor (hEGF) | Lonza | CC-4107 |
| Horse Serum | Gibco | 16050122 |
| Fetal Bovine Serum | Sigma | F2442 |
| Fetal Bovine Serum | Life Technologies | 10437028 |
| 0.25% Trypsin-EDTA (1X) | Gibco | 25200-056 |
| 0.05% Trypsin-EDTA (1X) | Gibco | 25300-054 |
| Penicillin Streptomycin (10,000 U/mL Pen, 10,000 ug/mL Strep) | Gibco | 15140-122 |
| Glycine | Sigma-Aldrich | G7126-100G |
| Tween 20, lot# SLBS9921 | Sigma-Aldrich | P9416-100mL |
| MEM Non-Essential Amino Acids (100X) | Gibco | 11140-050 |
| Bovine Serum Albumin (BSA), Protease-free powder | Fisher Scientific | BP9703-100 |
| PureCol, Purified Bovine Type I Atelo-Collagen Solution, 3 mg/mL | Advanced BioMatrix | 5005 |
| DPBS, no calcium, no magnesium | Gibco | 14190144 |
| PBS, pH 7.4 | Gibco | 10010023 |
| HBSS, no calcium, no magnesium, no phenol red | Life Technologies | 14175145 |
| Paraformaldehyde Solution, 4% in PBS | Thermo Scientific | J19943K2 |
| Triton X-100 | Acros Organics | A0376210 |
| QIAzol Lysis Reagent | Qiagen | 79306 |
| MitoTracker Deep Red | Thermo Fisher | M22426 |
| Dispase II (activity $\geq 0.5$ units/mg solid) | Sigma | D4693-1G |
| Collagenase Type II (654 U/mg, non-specific proteolytic activity 487 U/mg) | Life Technologies | 17101015 |
| Propidium Iodide - 1.0 mg/mL Solution in Water | Life Technologies | P3566 |
| Isofluorane | Vet One | 502017 |
| Corning CellTak | Fisher Scientific | C354240 |
| OCT | fisher scientific | 23-730-571 |
| ActinRed™ 555 ReadyProbes® | Invitrogen | R37112 |
| coverslips | Fisher scientific | S3023 |
| Dako fluorescent medium | Agilent | S3023 |
| Prolong Diamond | Invitrogen | P36970 |
| Direct Red | Fisher scientific | AAB2169306 |
| Picric Acid | Sigma | P6744-1GA |
| Glacial Acetic Acid | Fisher scientific | BP2401-500 |
| Hematoxylin | Fisher scientific | 3530-16 |
| Eosin | Millipore Sigma | 588X-75 |
| Safeclear | Fisher scientific | 23-044192 |
| PermOUNT | Fisher Scientific | SP15-100 |
| Goat Serum | Millipore Sigma | G9023 |

| Deposited Data |  |  |
| --- | --- | --- |
| Software and Algorithms |  |  |
| DESeq2 (v.1.26.0) | Love MI, Huber W, Anders S (2014) | https://www.genomebiology.biomedcentral.com/articles/10.1186/s13059-014-0550-8 |
| R v.4.0.0 | The R Foundation for Statistical Computing | https://www.r-project.org/ |
| WebGestaltR | Liao Y et al. (2019) | doi.org/10.1093/nar/gkz401 |
| Flexbar | Roehr et al. (2017) | [doi: 10.1093/bioinformatics/btx330](https://doi.org/10.1093/bioinformatics/btx330) |
| Kallisto (v.0.46.1) | Bray et al. (2016) | [doi:10.1038/nbt.3519](https://doi.org/10.1038/nbt.3519) |
| tximport (v.1.14.0) | Soneson C, Love MI, Robinson MD (2015) | doi.org/10.12688/f1000research.7563.2 |
| DP_GP_cluster | McDowell et al., (2018) | https://github.com/PrincetonUniversity/DP_GP_cluster |
| ComplexHeatmap | Gu Z, Eils R, Schlesner M (2016) | doi.org/10.1093/bioinformatics/btw313 |
| Escher | King ZA et al. (2015) | dx.doi.org/10.1371/journal.pcbi.1004321 |

### Myofiber Size and Quantification

Quadriceps muscles were mounted in OCT (Fisher Scientific # 23-730-571), snap frozen in liquid nitrogen, and sectioned in a cryotome at 10 μm thickness per section. Sections were stained with ActinRed™ 555 ReadyProbes^®^ (Invitrogen #R37112) by diluting two drops of the reagent in 1 mL of 1X PBS, WGA 488 (Biotium #29022) at 5 μg/ml concentration, and DAPI at 2 ug/mL concentration for one hour at room temperature. After the incubation period, dyes were washed three times for five minutes each in 1X PBS. Slides were mounted with Dako fluorescent medium (S3023) and coverslipped. Minimum Feret diameter distributions across whole sections were quantified using custom scripts in FIJI. Error bars for each bin were plotted by calculating the mean and standard error for three technical replicates and three biological replicates per bin.

### Fiber type stain and Quantification

Quadriceps muscles were mounted in OCT (Fisher Scientific # 23-730-571), snap frozen in liquid nitrogen, and sectioned in a cryotome at 10 μm thickness for each section. Slides were first rinsed in PBS three times for 5 minutes with gentle shaking followed by a 5-minute incubation with 0.2% Triton X-100 (Acros Organics #A0376210) in 1X PBS for 5 minutes and rinsed again in 1X PBS as before. Tissue sections were then blocked in M.O.M. mouse IgG blocking reagent (Vector Laboratories #BMK-2202) overnight in a hydrated chamber. Blocking reagent was then washed three times for 5 minutes in 1X PBS, followed by a 5-minute incubation in M.O.M. diluent. Primary antibodies were incubated at 4°C overnight in a hydrated chamber and diluted in M.O.M. diluent as follows: BA-D5 IgG2b (DSHB University of Iowa) at 1:100 dilution, SC-71 (DSHB University of Iowa) at 1:500 dilution, and BF-F3 (DSHB University of Iowa) at 1:100 dilution. After three 5-minute 1X PBS washes, secondary antibodies and WGA were incubated for one hour at room temperature at the following concentrations: Alexa Fluor 647 Goat anti-Mouse IgG2b (Invitrogen #A-21242) at 1:300 dilution, Alexa Fluor 488 Goat anti-Mouse IgG1 (Invitrogen #A-21121) at 1:300 dilution, Alexa Fluor 555 Goat anti-Mouse IgM (Invitrogen #A-21426) at 1:300 dilution, and WGA 405 (Biotium #29027) at 100 μg/ ml concentration. Secondary antibodies and WGA were washed three times for 5 minutes in 1X PBS. Tissue sections were mounted with Dako fluorescent medium (Agilent #S3023) and coverslipped. Whole section images were taken using a Nikon A1 confocal microscope. Fiber type distributions of quadriceps whole section were manually quantified in FIJI. Statistical significance was calculated by performing a two-sided Mann-Whitney *U*-tests across three biological replicates with three technical replicates per sample group.

### Pax7/Phospho-S6 Staining and Quantification

Quadriceps and TA muscles were mounted in OCT, snap frozen in liquid nitrogen, and sectioned in a cryotome at 10 μm thickness for each section. Tissue sections were fixed with 4% PFA (Fisher Scientific # AAJ19943K2) for 10 minutes and washed three times for 5 minutes each in 1X PBS. Endogenous peroxidases were blocked with 100X H2O2 for 10 minutes at room temperature from the Tyramide SuperBoost Goat anti-Mouse AF555 or AF488 kit (Invitrogen #B40913, #B40912) followed by 1X PBS washes. Slides were incubated in citrate buffer at room temperature for 2 minutes, placed in citrate buffer at 65°C and heated to 92°C. Once at 92°C, tissue sections were incubated for 11 minutes and brought back to room temperature. Sections were then blocked in M.O.M. mouse IgG blocking reagent (Vector Laboratories #BMK-2202) and 10% goat serum (Millipore Sigma#G9023) for one hour at room temperature followed by three 5-minute 1X PBS washes. Sections were incubated with Pax7 from supernatant (DSHB University of Iowa), and anti-rabbit Phospho-S6 (Cell Signaling #4858) overnight at 4°C in a humidified chamber in M.O.M. diluent at 1:10 and in 10% goat serum at 1:50 dilutions, respectively. Primary antibodies were washed three times for 5 minutes in 1X PBS and secondary goat anti-mouse Poly HRP antibody from Tyramide SuperBoost Goat anti-Mouse AF555 or AF488 kits were added for 1 hour at room temperature, followed by 1X PBS washes. Tyramide working solution was then incubated for 10 minutes, followed by 1X PBS washes. WGA 488 (Biotium #29022) at 50 μg/ml concentration, DAPI at 2μg/ml concentration, and Alexa Fluor 647 or Alexa fluor 555 Goat anti-rabbit (Invitrogen #A-21244, #A-21428) were incubated for 1 hour at room temperature. Slides were then mounted with Dako fluorescent media, and coverslipped (S3023). Whole section images were taken using a Nikon A1 confocal microscope. Satellite cells positive and negative for phospho-S6 were counted by hand using FIJI. Statistical significance was calculated by unpaired two-sided Student’s *t*-test across three technical replicates and three biological replicates per sample group.

### p62 Staining and Quantification

TA muscles were mounted in OCT, snap frozen, and sectioned in a cryotome at 10 μm thickness. Tissue sections were re-hydrated with three washes of 1X PBS of 5 minutes each. Sections were treated with 0.2% Triton X-100 in 1X PBS for 5 minutes followed by three washes of 1X PBS. Sections were blocked with 0.1% goat serum (Millipore Sigma#G0923) for 1 hour at room temperature followed by overnight primary antibodies incubation (SQSTM1/p62 Rabbit mAb #23214) at 1:200 dilution in 0.1 % goat serum in 4°C. Primary antibodies were washed 3 times with 1X PBS for 5 minutes each followed by incubation with goat anti-rabbit Alexa Fluor 555 (Invitrogen# A27039) 1:300 dilution in 0.1 % goat serum), WGA 488 (100 μg/ml, Biotium #29022), and DAPI at 2 μg/ml for 1 hour at room temperature. Slides were then mounted with Prolong Diamond and coverslipped (Invitrogen #P36970). Whole section images were taken using a Nikon A1 confocal microscope. FIJI was used to quantify the total number of p62- positive fibers and minimum Feret diameter.

### Picrosirius Stain

Tissue sections were fixed in 4% PFA (Fisher Scientific # AAJ19943K2) for 15 minutes at room temperature. 4% PFA was washed two times with 1XPBS and 2 times with deionized water for five minutes each. Sections were air dried for 15 minutes at room temperature. Sections were incubated for 1 hr. with Sirius Red dye (Fisher scientific #AAB2169306, Sigma #P6744-1GA) in a humidifier chamber. Sirius red dye was washed two times for 5 minutes each with acidified water (Fisher Scientific BP2401-500) and deionized water at room temperature. Sections were dehydrated by quick immersions in 50%, 70%, 70%, 90%, 100%, and 100% ethanol series followed by two 5 minutes incubations at room temperature. Coverslips were mounted with permount (Fisher Scientific #SP15-100). Whole section images were taken using a motorized Olympus IX83 microscope

### Hematoxylin and Eosin stain

Slides were submerged in hematoxylin (fisher scientific # 3530-16) for two minutes followed by two sets of ten quick immersions in distilled water. Slides were then submerged in Scott’s tap water and distilled water for one minute as well as one set of ten immersions in 80% ethanol. After one minute of Eosin (Millipore Sigma # 588X-75) immersion slides were immersed ten times in two different 95% ethanol baths and a 100% ethanol bath. Finally, slides were submerged in safeclear (fisher scientific #23-044192) for one minute, and two drops of permount (Fisher Scientific #SP15-100) was added before placing the coverslip on top. Whole section images were taken using a motorized Olympus IX83 microscope.

### Satellite Cell Isolation via Fluorescence-Activated Cell Sorting.

For tissue collection, mice were anesthetized with 3% isoflurane, then euthanized by cervical dislocation, bilateral pneumothorax and removal of the heart. Hind limb muscles (tibialis anterior, gastrocnemius, and quadriceps) of wild type and sestrin double knockout mice were quickly harvested using sterile surgical tools and placed in separate plastic petri dishes containing cold PBS. Using surgical scissors, muscle tissues were minced and transferred into 50 mL conical tubes containing 20 mL of digest solution (2.5 U/mL Dispase II and 0.2% [~5,500 U/mL] Collagenase Type II in DMEM media per mouse). Samples were incubated on a rocker placed in a 37°C incubator for 90 min with manual pipetting the solution up and down to break up tissue every 30 minutes using an FBS coated 10 mL serological pipette. Once the digestion was completed, 20 mL of F10 media containing 20% heat inactivated FBS was added into each sample to inactivate enzyme activity. The solution was then filtered through a 70 μm cell strainer into a new 50 mL conical tube and centrifuged again at 350xg for 5 min. The pellets were resuspended in 6 mL of staining media (2% heat inactivated FBS in Hank’s Buffered Salt Solution - HBSS) and divided into separate FACS tubes. The FACS tubes were centrifuged at 350xg for 5 min and supernatants discarded. The cell pellets were then re-suspended in 200 μL of staining media and antibody cocktail containing Sca-1:APC (1:400), CD45:APC (1:400), CD11b:APC (1:400), Ter119:APC (1:400), CD29/B1-integrin:PE (1:200), and CD184/CXCR-4: BIOTIN (1:100) and incubated for 30 minutes on ice in the dark. Cells and antibodies were diluted in 3mL of staining solution, centrifuged at 350xg for 5 min, and supernatants discarded. Pellets were resuspended in 200uL staining solution containing PECy7: STREPTAVIDIN (1:100) and incubated on ice for 20 minutes in the dark. Again, samples were diluted in 3mL staining solution, centrifuged, supernatants discarded, and pellets re-suspended in 200uL staining buffer. Live cells were sorted from the suspension via addition of 1 μg of propidium iodide (PI) stain into each experimental sample and all samples were filtered through 35 μm cell strainers before the FACS. Cell sorting was done using a BD FACSAria III Cell Sorter (BD Biosciences, San Jose, CA) and APC negative, PE/PECy7 double-positive MuSCs were sorted into staining solution for immediate processing.

### Preparation of RNA-Seq Libraries and Sequencing

MuSCs were FACS sorted directly into Trizol and snap frozen. Samples were subsequently thawed, and RNA was extracted using a Qiagen miRNeasy Micro Kit as per manufacturer’s instructions. The integrity of the isolated RNA was verified with a Bioanalyzer (Agilent 2100) and 1-10 ng of high-quality RNA (RIN>8) was used to produce cDNA libraries using the SmartSeq v4 protocol (Clontech) as per the manufacturer’s instructions. cDNAs were prepared into sequencing libraries using 150 pg of full-length cDNA amplicons (Nextera XT DNA Library Preparation Kit, Illumina) with dual index barcodes. Barcoded cDNA libraries were pooled into a single tube and sequenced on a NextSeq (Illumina) using 76-bp single-ended reads.

### RNA-Seq Data Processing and Analysis

#### Gene Expression Estimation

Single-end RNA-Seq data were trimmed using Flexbar (v3.5.0) and pseudo-aligned to the mouse reference genome (GRCm38.p6) using Kallisto (v.0.46.1). Reads averaged 41.75M per sample. The full Kallisto command was as follows:

kallisto quant -b 100 --single -1 300 -s 30 -i **[mm10.idx]** -t 45 -o **[output folder] [trimmed FASTQ]**

#### Differential Gene Expression

The estimated transcript abundances were summarized to gene-level count matrices using tximport and genes containing at least one read were retained. Differentially expressed genes in treated (KO) samples relative to untreated controls (WT) at each timepoint (days=0,7,21) were identified using DESeq2 in R with a design formula: *Count ~ group,* with group={day+treatment}, day={0,7,21}, and treatment={WT,KO}. Surrogate variable analysis was performed on the rlog-transformed count matrix using the SVA package with a null model of *rlog(Count) ~ 1* and a design matrix of *rlog(Counts) ~ group.* Contributions from the surrogate variable were quantified and removed from the rlog-transformed count matrix for downstream analyses. Pairwise Spearman correlation analysis was performed between all replicates and replicates that had r < 0.9 with other replicates were excluded from further analysis. Pairwise contrasts were examined to find differentially expressed genes between SKO vs WT on each day post injury. Differentially expressed genes were selected using a false discovery rate (FDR) cutoff of 0.01 and a Log_2_ fold change cutoff of 1.

#### Time-series Clustering of Differential Genes

Genes that were differentially expressed on at least one day post injury were pooled and submitted for time-clustering using the Dirichlet Process Gaussian Process (DPGP) algorithm. To prepare inputs for the algorithm, regularized log-transformed counts were averaged across biological replicates on each day post injury and the fold change in averaged counts was calculated between SKO vs WT. The resulting fold changes for each gene were standardized across time points using a z-score transformation. DPGP clustering was performed using the default parameters. The full command was as follows:

DP_GP_cluster.py -i **[fold change z-scores]** -o ./**[output file prefix]**

DPGP assigned each gene a unique time-dependent cluster based on similar expression dynamics, and clusters that exhibited similar temporal dynamics were manually combined into a single cluster. Log-fold z-scores were plotted as a function of time for each cluster and a heatmap of differentially expressed genes grouped by DPGP cluster was plotted using the ComplexHeatmap package in R.

#### Pathway Enrichment Analysis

The top 100 upregulated and downregulated genes for 0 days post injury were submitted for Gene Ontology (GO) term and Kyoto Encyclopedia of Genes and Genomes (KEGG) pathway over-representation analysis using the WebGestaltR package in R. A GO Biological Process (GO:BP) reference curated to remove redundant terms was used. Only KEGG and GO:BP terms containing between 10 and 500 genes were considered for enrichment. Genes in each DPGP cluster analyzed using the same procedure. Redundancy in the enriched terms (FDR < 0.05) was reduced using the Affinity Propagation or Weighted Set Cover algorithms in WebGestaltR. Percent coverage of all terms produced by the Weighted Set Cover algorithm was at least 95%. The resulting terms were plotted using the ggplot2 package and ComplexHeatmap packages in R.

#### Derivation of reaction fluxes using gene expression data and genome-scale metabolic modeling

Gene expression data from SKO and WT MuSCs were normalized (z-scores) across each gene. Normalized data were compared using an unpaired two-sided t-test in MATLAB R2018b to identify differentially expressed genes (p<0.05, |z|>1.5) between SKO and WT MuSCs before injury. Differentially expressed genes for SKO and WT MuSCs were used as input to derive reaction flux information from a human genome-scale metabolic model (RECON1)^30,45^ that maximizes and minimizes flux through metabolic reactions associated with upregulated and downregulated genes, respectively, based on internal gene-protein-reaction annotations. Reaction flux data were generated using a linear optimization version of the iMAT algorithm^30,45^ with the RECON1 model and the following parameters: (rho = 1E-3, kappa = 1E-3, epsilon = 1, mode = 0). Reaction fluxes in SKO MuSCs were then normalized against those in WT MuSCs and normalized (z-scores) using R. Normalized flux differences were overlaid onto maps of metabolic pathways using Escher^46^ for visualization. Reaction flux differences with |z|>3 were considered significant.

## Competing interests

The authors declare no competing interests.

## Notes

https://www.ncbi.nlm.nih.gov/geo/query/acc.cgi?acc=GSE162191

### Competing Interest Statement

The authors have declared no competing interest.

