## Supplemental figure 1 for "Sestrins regulate age-induced deterioration of muscle stem cell homeostasis"

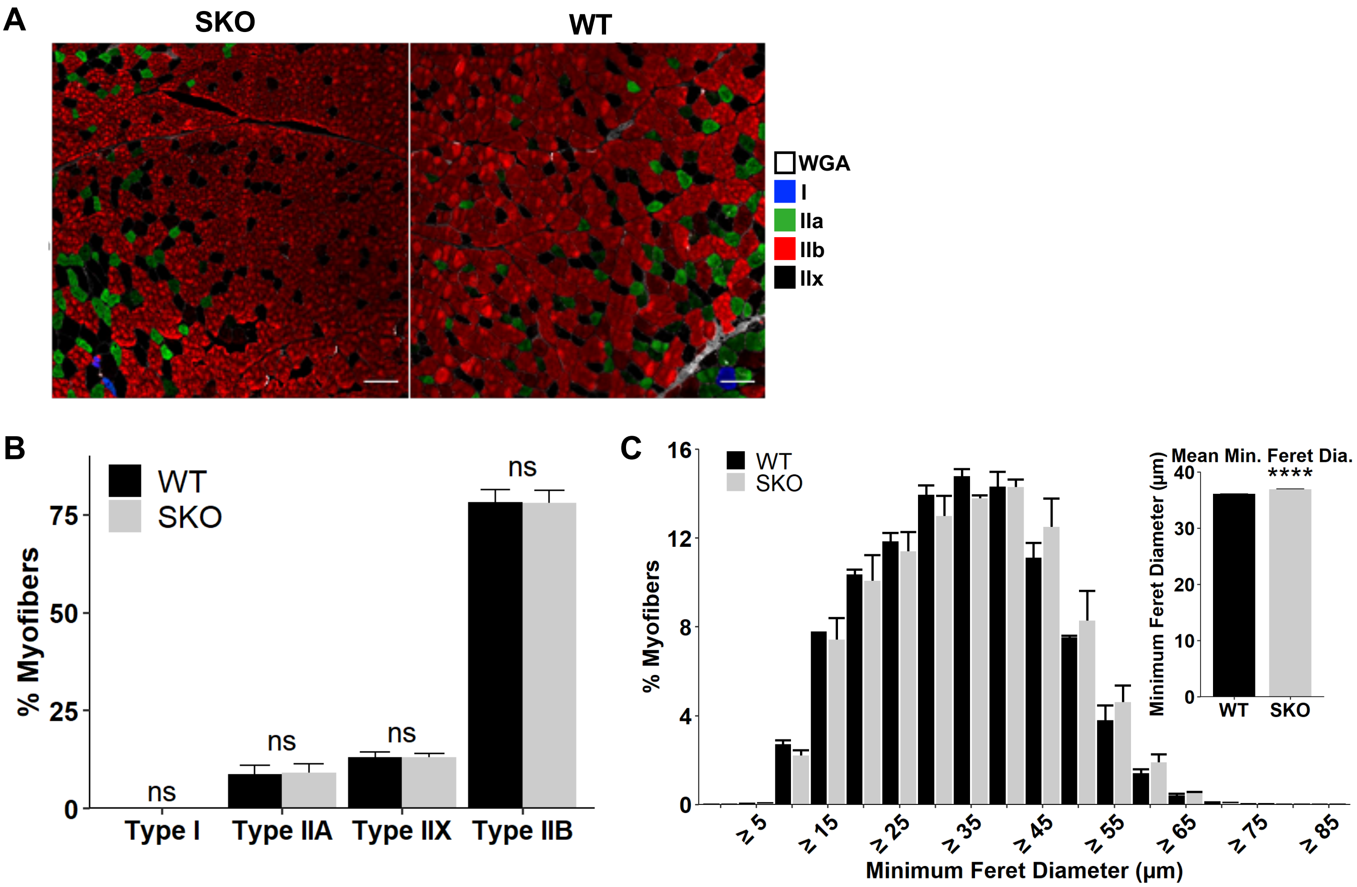


**Supplementary Figure 1. Related to Figure 1. A)** Representative immunohistochemical images of fiber-type analyses in whole quadriceps muscle sections stained with antibodies against Type I (blue), Type IIa (green), and Type IIb (red) fibers. Type IIx fibers lack staining. Connective tissue was stained with wheat germ agglutinin (WGA) (white). Scale bars are 100 μm. **B)** Distribution of fiber types in the rectus femoris of the quadriceps muscle. (WT: n=3, SKO: n=3). Statistical comparisons are two-sided Mann-Whitney *U*-tests. **C)** Size distributions of WT and SKO myofibers across whole quadriceps muscle sections. Statistical comparisons are two-sided unpaired Student's *t*-tests with Holm multiple testing correction for the histogram. (WT: n=3, SKO: n=3). Inset is mean myofiber size per condition. All data are shown as mean ± SEM. Statistical significance thresholds were set at $p<0.05$ and $p_{adj}<0.05$. Unmarked comparisons lack statistical significance.

**SUPP. FIGURE 1:**
