## Supplemental Figure 2 for "Sestrins regulate age-induced deterioration of muscle stem cell homeostasis"

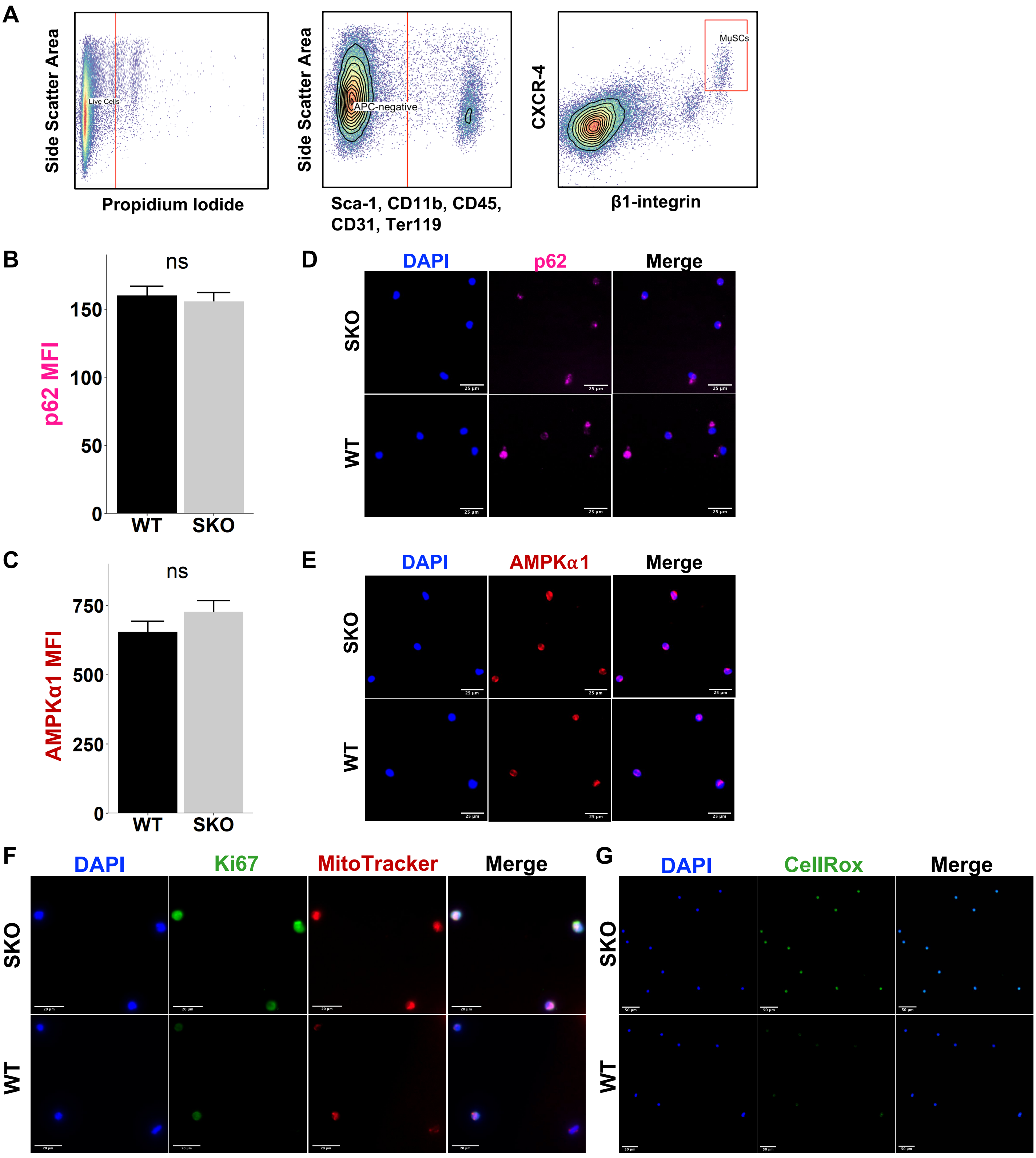


**Supplementary Figure 2. Related to Figure 1. A)** Representative muscle stem cell (MuSC) isolation plots using FACS gating for Sca-1$^{-}$, CD45$^{-}$, CD11b$^{-}$, Ter-119$^{-}$, **β**1-integrin$^{+}$, and CXCR-4$^{+}$. Mean fluorescence intensity (MFI) of **B)** p62 and **C)** AMPKα1 in SKO and WT MuSCs fixed immediately after isolation. Representative DAPI-counterstained immunofluorescence images of **D)** p62 and **E)** AMPKα1 in MuSCs after 3 days of culture. Scale bars are 25 µm. MFI of **F)** Immunofluorescence staining for Ki67 (green), MitoTracker (red), and DAPI (blue) in SKO and WT MuSCs fixed immediately after isolation. Scale bars are 20 µm. **G)** Immunofluorescence staining for CellRox (green) and DAPI (blue) in SKO and WT MuSCs fixed immediately after isolation. Scale bars are 50 µm. All data are shown as mean ± SEM. Statistical comparisons are two-sided unpaired Student's *t*-tests.

**SUPP. FIGURE 2**
