## Supplemental Figure 3 for "Sestrins regulate age-induced deterioration of muscle stem cell homeostasis"

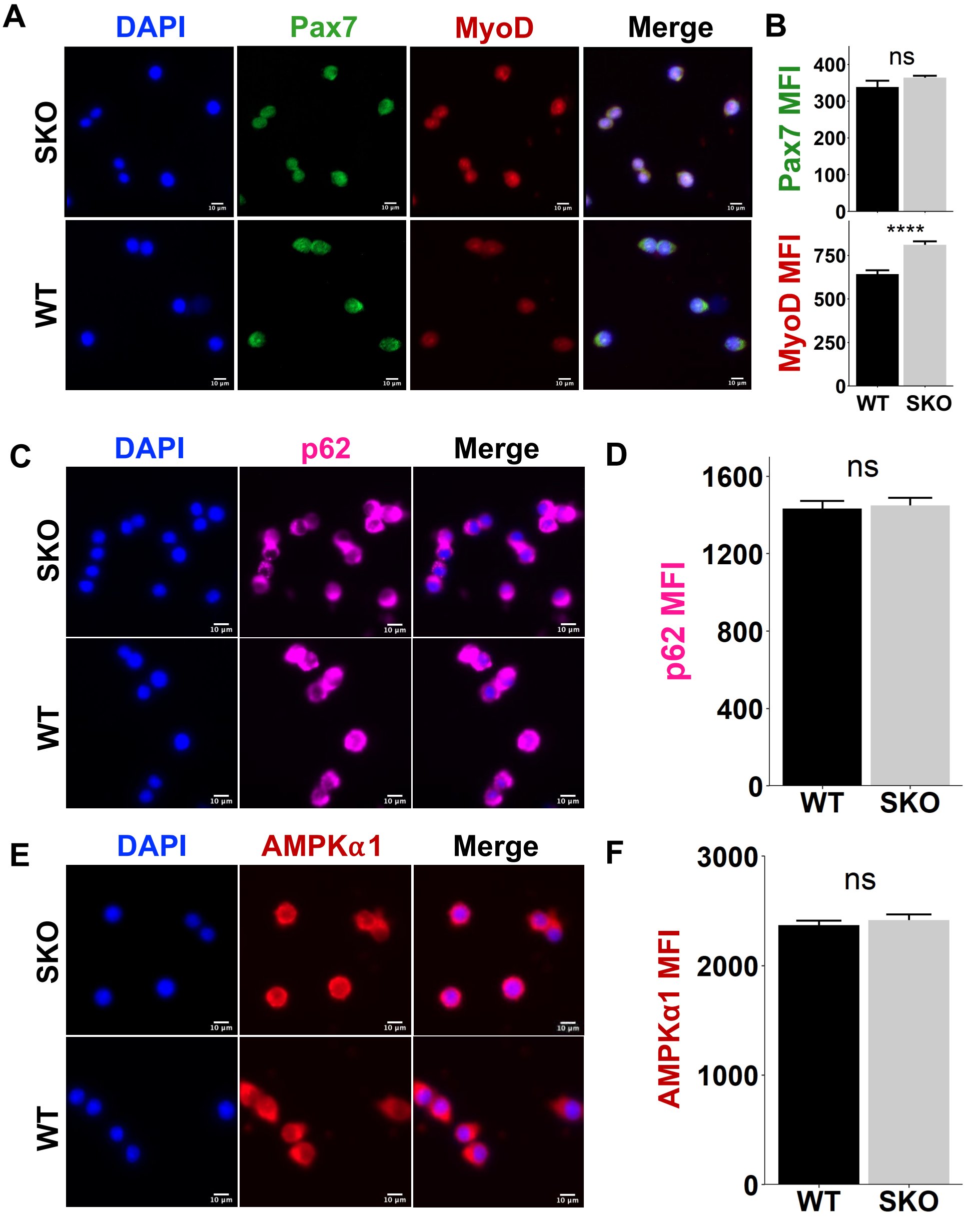


**Supplementary Figure 3. Related to Figure 1. A)** Immunofluorescence staining for Pax7 (green), MyoD (red) and DAPI (blue) in SKO and WT MuSCs after 3 days of culture in activating conditions. **B)** Mean fluorescence intensity (MFI) of Pax7 and MyoD in SKO and WT MuSCs after 3 days of culture. Immunofluorescence staining for **C)** p62 and **E)** AMPKα1 in MuSCs after 3 days of culture. Quantification of MFI from **D)** p62 and **F)** AMPKα1 in SKO and WT MuSCs fixed after 3 days of culture in activating conditions. All scale bars are 10 µm. All data are shown as mean ± SEM. Statistical comparisons are two-sided unpaired Student's *t*-tests. ****p<0.0001.

SUPP. FIGURE 3
