## Supplemental Figure 4 for "Sestrins regulate age-induced deterioration of muscle stem cell homeostasis"

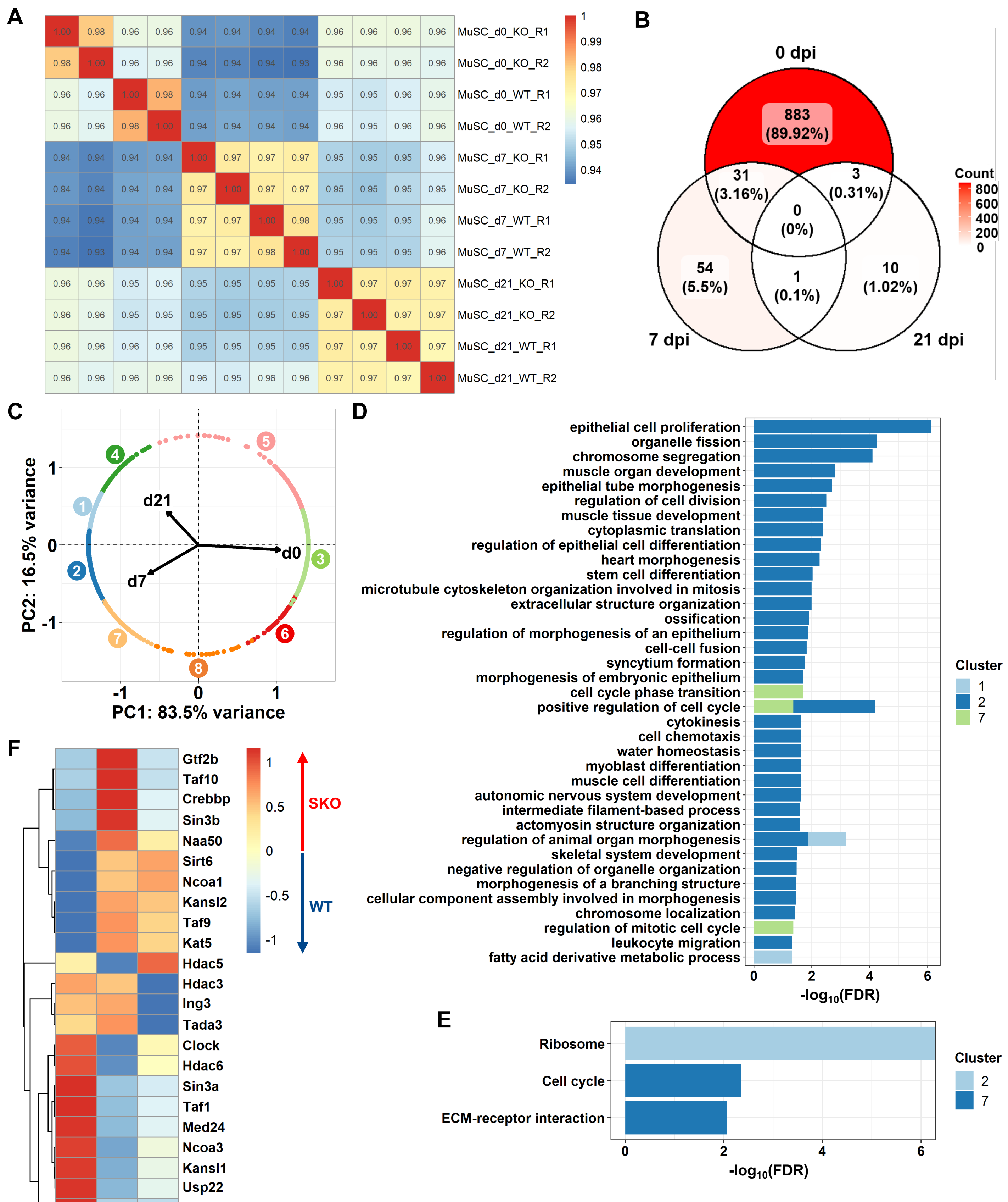

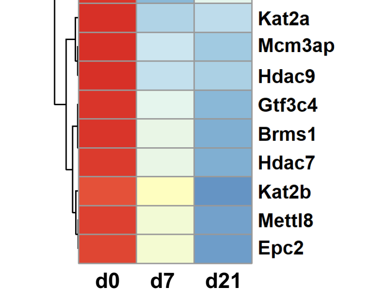

**Supplementary Figure 4. Related to Figure 2. A)** Spearman correlation heatmap between all biological replicates. **B)** Venn diagram of differentially expressed genes (padj<0.01, $Log_2$ Fold Change>1) in freshly isolated MuSCs at 0, 7, and 21 days post injury (dpi). **C)** Principal component analysis (PCA) plot of differentially expressed genes colored by DPGP cluster with loadings of dpi. **D)** Stacked bar plot of enriched GO:BP terms (FDR<0.05) per DPGP cluster from WebGestaltR analysis. **E)** Bar plot of enriched KEGG terms (FDR<0.05) per DPGP cluster from WebGestaltR analysis. **F)** Standardized heatmap (z-scores) of differentially expressed genes (padj<0.05, $Log_2$ Fold Change>0.26) from WT and SKO MuSCs in GO terms for histone deacetylase (GO:0004407) or acetyltransferase (GO:0004402) activity at 0, 7, and 21 dpi.

**SUPP. FIGURE 4**
