## Supplemental Figure 5 for "Sestrins regulate age-induced deterioration of muscle stem cell homeostasis"

A

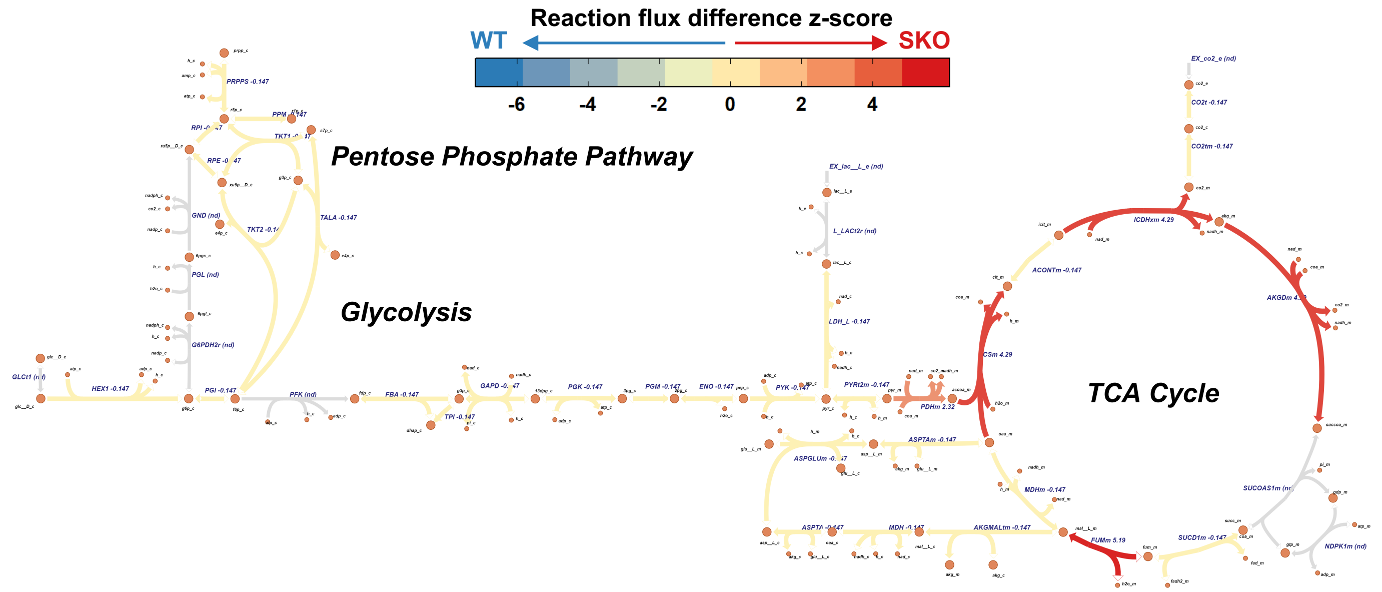


B

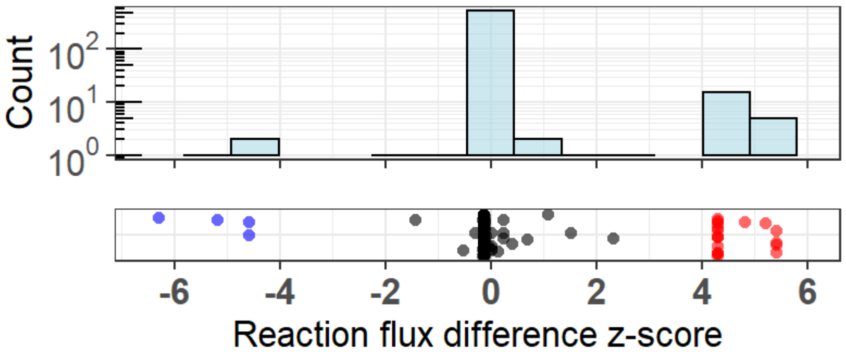


C

| ID | Name | Subsystem |
|---|---|---|
| ACSm | Acetyl CoA synthetase | Glycolysis / Gluconeogenesis |
| CSm | Citrate synthase | Citric Acid Cycle |
| GLYATm | Glycine C-acetyltransferase | Glycine, Serine, and Threonine Metabolism |

D

| ID | Name | Subsystem |
|---|---|---|
| GLUDxm | Glutamate dehydrogenase, mitochondria | Glutamate metabolism |
| RADH | Retinal dehydrogenase | Vitamin A metabolism |
| ACACtx | Hydrogen peroxide transport via diffusion | Transport, Extracellular |

**Supplementary Figure 5. Related to Figure 2. A)** Metabolic flux map of glycolysis, the pentose phosphate pathway, and the TCA cycle colored by RECON1 reaction flux difference z-scores between SKO and WT muscle stem cells (MuSCs). **B)** Histogram (top) and jitter plot (bottom) of reaction flux difference z-scores. Reactions with flux difference z-scores beyond the |z|>3 threshold are colored by z-score polarity (red, SKO; blue, WT). Representative significant fluxes for each metabolic subsystem in SKO **(C)** and WT **(D)** MuSCs are shown.

**SUPP. FIGURE 5**
