## Supplementary Figure 6 for "Sestrins regulate age-induced deterioration of muscle stem cell homeostasis"

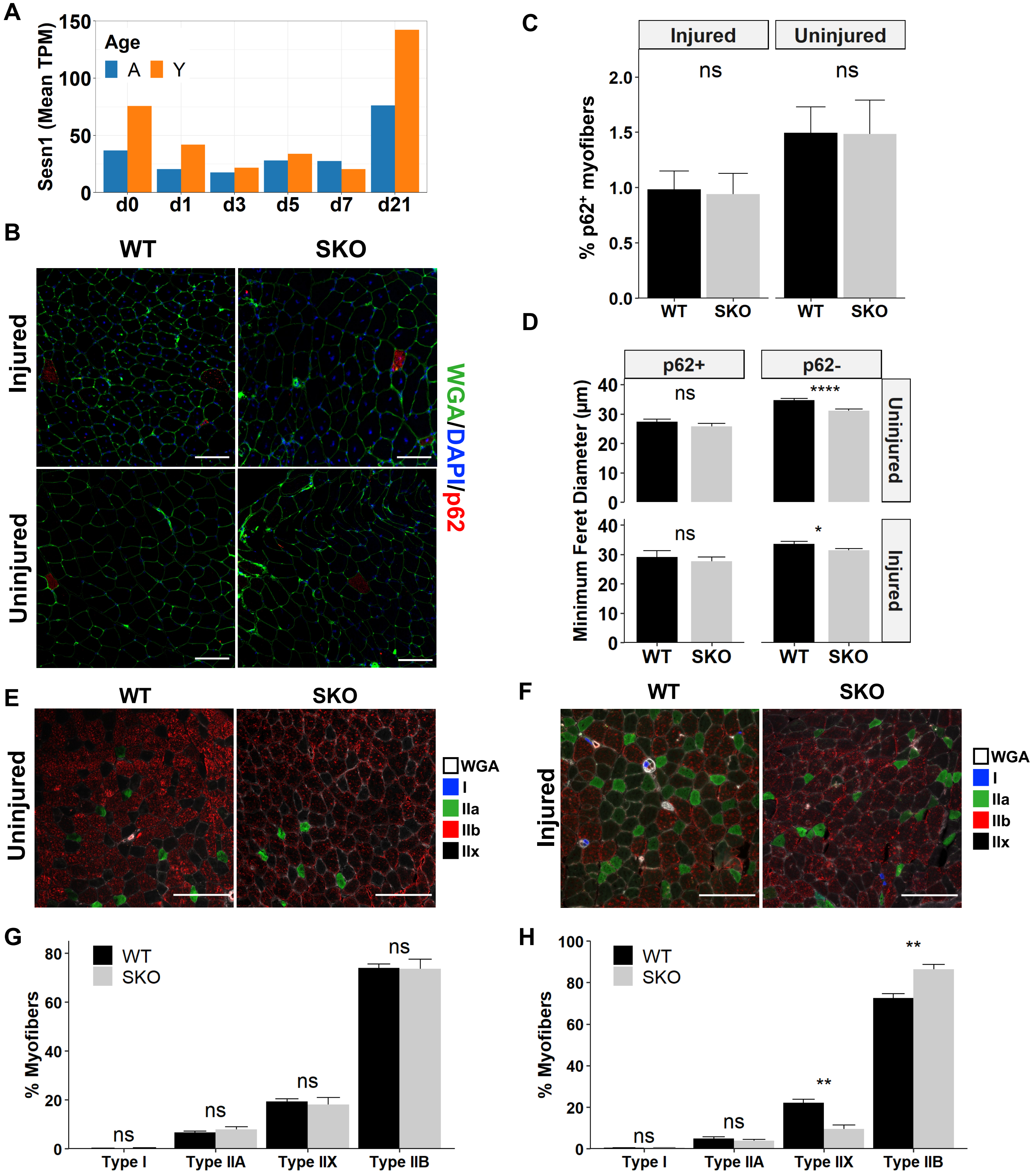


**Supplementary Figure 6. Related to Figures 3 and 4. A)** Average Sesn1 expression in young (2-3 months) and aged (22-24 months) MuSCs at 0, 1, 3, 5, 7, and 21 days after $BaCl_2$ hindlimb (gastrocnemius and tibialis anterior) injury. **B)** Representative immunohistochemical images of p62 in whole TA muscle sections. Scale bars 100 µm. **C)** Percentage of $p62^+$ myofibers in whole TA muscle sections from WT and SKO mice. **D)** Minimum feret diameters of $p62^+$ and $p62^-$ myofibers in injured and uninjured whole TA muscle sections from WT and SKO mice. Statistical comparisons are two-sided unpaired Student's *t*-tests. Representative immunohistochemical images of fiber-type analyses in **E)** uninjured and **F)** injured whole TA muscle sections from WT and SKO mice. Sections were stained with antibodies against Type I (blue), Type IIa (green), and Type IIb (red) fibers. Type IIx fibers lack staining. Connective tissue was stained with wheat germ agglutinin (WGA) (white). Scale bars are 100 µm. Distributions of fiber types in whole TA muscle sections of **G)** uninjured and **H)** injured mice. Statistical comparisons are two-sided U-tests. All data are shown as mean ± SEM (Injured – WT: n=4, SKO: n=3; Uninjured – WT: n=5, SKO: n=3). *p<0.05, **p<0.01, and ****p<0.0001.

**SUPP. FIGURE 6**
